# Rapid speciation in small populations challenges the dominance of speciation by local adaptation

**DOI:** 10.64898/2026.02.19.706750

**Authors:** Pierre Veron, Anaïs Spire, Agathe Chave-Lucas, Tatiana Giraud, Hélène Morlon

**Affiliations:** Institut de Biologie de l’École Normale Supérieure, CNRS, INSERM, Université PSL, Paris, France; Écologie Société et Évolution, CNRS, Université Paris-Saclay, AgroParisTech, Gif-sur-Yvette, France

**Keywords:** speciation, micro–macro, holey adaptive landscape, plants

## Abstract

Speciation — the process by which two lineages become reproductively isolated — plays a key role in the emergence and maintenance of biodiversity. Yet, our understanding of the time it takes for speciation to occur, and of the microevolutionary processes that influence this tempo, remains limited. Here, we thoroughly characterize how population size, mutation rate, local adaptation and migration are expected to influence the duration of speciation, as well as the shape of the “grey zone” of speciation. We show that the relationship between population size and speciation duration is indicative of the speciation mode, as faster speciation in smaller populations is the signature of speciation by neutral divergence. Leveraging genomic estimates of population size and speciation duration across 196 pairs of plant species, we uncover a positive association between population size and speciation duration. Taken together, these results challenge the view that speciation by divergent local adaptation is the source of much of species diversity.

## 1 Introduction

Speciation — the process by which a new species emerge — is one of the most fundamental processes in biology. It is at the origin of species diversity. In spite of substantial research on speciation, many controversies remain on its mode, i.e, how it occurs, and tempo, i.e., at which pace it occurs (Coyne and Orr 2004). There is however a general consensus that speciation takes time (Benton and Pearson 2001; Etienne et al. 2014), except in exceptional cases, such as speciation by polyploidization in plants (Rieseberg and Willis 2007) or speciation by host shifts in pathogens (Giraud et al. 2010). Intuitively, fast-speciating groups will experience speciation events more frequently than slow-speciating ones. Together with the frequency of extinction events, these differences can explain differences in species richness across groups, and together with the frequency of dispersal events, differences in species richness across geographic regions (Wiens and Donoghue 2004; Schluter and Pennell 2017).

Insights on the tempo of speciation can be gained by fitting diversification models to extant phylogenies (Nee 2006; Morlon et al. 2024). Fitting birth-death models representing lineages birth (speciation) and death (extinction), assumed to occur instantaneously, provides estimates of speciation and extinction rates, i.e., estimates of the average number of events occurring per lineage in a given amount of time. Such studies have shown that speciation rates vary by several orders of magnitude across lineages (Rabosky 2016; Maliet et al. 2019; Quintero et al. 2024). For example, estimates range from 0.01 to 5 spp Myr*^−^*^1^ in birds (Maliet et al. 2019), and from 0.005 to 1.5 spp Myr*^−^*^1^ in mammals (Quintero et al. 2024).

A central challenge lies in understanding if and how microevolutionary (intraspecific) processes, such as drift, selection and migration, can explain this speciation rate heterogeneity (Harvey et al. 2019; Rolland et al. 2023; Morlon et al. 2024). In comparison with the numerous studies that have investigated correlates of speciation rates with species traits, as well as the abiotic and biotic environment they experience (Benton and Pearson 2001; Lagomarsino et al. 2016; Schluter and Pennell 2017; Landis et al. 2022; Wiens 2024), empirical studies correlating speciation rates to metrics reflecting microevolutionary processes remain rare. The few studies that have investigated the relationship between speciation rates and such parameters have found contrasting or counter-intuitive results, in particular in terms of correlations with the velocity of reproductive isolation (Rabosky and Matute 2013), population divergence rates (Harvey et al. 2017; Harvey et al. 2019), population structure (Singhal et al. 2018; Burbrink et al. 2023), substitution rates (Lanfear et al. 2010; Goldie et al. 2011), or genetic diversity (Huang et al. 2018; Perez-Lamarque et al. 2022; Afonso Silva et al. 2025).

Mathematical models have been central in our understanding of the dynamics of speciation, and of the microevolutionary processes that influence these dynamics (Gavrilets 2003; Coyne and Orr 2004). In particular, Gavrilets’ holey adaptive landscape (HAL) provided a mathematically tractable extension of Wright’s rugged adaptive landscape (Wright 1932) to high dimensional genotypic spaces (Gavrilets et al. 1998). He showed that — simply as a result of the high dimensionality of adaptive landscapes — populations can become separated by regions of low fitness (“holes”), and thus become reproductively isolated, without crossing deep valleys of low fitness by evolving along nearly neutral ridges (Gavrilets 2003). This framework is therefore well suited for analyzing how drift and selection jointly influence the dynamics of speciation. In his 1999 paper, Gavrilets derived a series of mathematical results describing the dynamics of divergence within- and between-populations under strict allopatry, in the presence of migration, and with local adaptation. These provide the foundation for obtaining analytical results concerning the expected duration of speciation, although those results were not derived. A later review focused on this question (Gavrilets 2003) but without taking into account within-population genetic variation. Yet, polymorphism can substantially alter the speed at which mutations causing reproductive incompatibilities accumulate, and therefore speciation duration (Gavrilets 1999).

Population size has occupied a central position in early debates about the relative role of drift and selection in the speciation process (Coyne and Orr 2004). This idea traces back to Mayr’s verbal “genetic revolution” hypothesis that population bottlenecks, for example during founder events, can trigger a shift in allele frequency over several linked loci leading to speciation (Mayr 1963). Under this “drift” hypothesis, speciation should be faster in smaller populations. Opponents of this hypothesis, however, predicted that speciation should be faster in larger populations, where natural selection is more efficient, facilitating divergent adaptation (Orr and Orr 1996). The relationship between population size and speciation duration could thus be indicative of the preponderance of speciation by local adaptation, which can occur with or without gene flow (Rundle and Nosil 2005). Indeed, the extent to which divergent adaptation is responsible for the accumulation of reproductive barriers remains highly debated (Schluter 2009; Anderson and Weir 2022).

While seemingly straightforward, the potential to test whether speciation occurs by local adaptation provided by the relationship between population size and speciation duration has not been widely used (but see Maya-Lastra and Eaton 2021). This is likely due to the absence of theoretical predictions for the duration of speciation as a function of population size, and to the difficulty of estimating both the duration of speciation and population sizes when speciation occurred. Recent developments in phylogenetic models of diversification, combined with the availability of large empirical datasets, have allowed for new tests of the relationship between proxies of population size, such as range size or nucleotide diversity, and speciation rates (Maya-Lastra and Eaton 2021; Smyčka et al. 2023; Afonso Silva et al. 2025). These tests, however, treat speciation as an instantaneous event, which rate is only weakly influenced by the duration of speciation (Veron et al. 2025), and can be confounded by the reciprocal effect of speciation on population sizes (Smyčka et al. 2023). Inferences from speciation genomics, on the other hand, can provide estimates of speciation duration *per se*, along with estimates of population sizes before and after the speciation event (Fraïsse et al. 2021).

Here, we begin by deriving new analytical predictions for the duration of speciation under the joint influence of drift, selection and gene flow, focusing on the effect of population size. Our derivations build upon Gavrilets’ holey adaptive landscape (HAL) model (Gavrilets 1999). We then design a test based on our theoretical results on the relationship between speciation duration and population size to assess speciation mode based on genomic data, which we apply to 196 pairs of plant species (Monnet et al. 2025).

## 2 Results and discussion

### Modelling the accumulation of reproductive incompatibilities between populations

We consider the HAL model of speciation (Gavrilets et al. 1998; Gavrilets 1999; Materials and methods). Initially, mutations arise in a population made of *N* individuals, with a per-individual per-generation mutation rate *ν*. Any pair of individuals is thus characterized by a genetic distance *d*, defined as the number of nucleotide sites at which they differ. If *d* is above a given threshold *K* called “genetic incompatibility threshold”, the two individuals do not interbreed, as a result of either post- or prezygotic incompatibilities (e.g., hybrid inviability or assortative mating). Hence *K* is the maximum number of loci on which two individuals can be different and still have a chance to mate successfully. This formulation is common to multiple speciation models (Higgs and Derrida 1992; de Aguiar et al. 2009; de Aguiar 2017; Hissa et al. 2026), motivated by the decrease in the reproductive success of two individuals with the genetic distance that separates them (Price and Bouvier 2002; Rabosky and Matute 2013; Christie and Strauss 2018; Christie et al. 2022); it simply reflects the presence of an isolation mechanism that prevents reproduction between genetically distant individuals. This formulation provides a mathematically tractable way to represent high dimensional, correlated adaptive landscapes (Gavrilets 1997, figure 1A), and to account for the multi-genic, epistatic, origin of incompatibilities. This model focuses on the consequence of incompatibilities — reproductive isolation — rather than on their nature (e.g., trait-based mate choice, Dobzhansky-Muller incompatibilities, DMIs; Dobzhansky 1936; Muller 1942). The specific form of the relationship between genetic distance and reproductive success has been highly debated (Matute et al. 2010; Städler et al. 2011). We can expect that it varies depending on the nature of incompatibilities between individuals (e.g., extrinsic *versus* intrinsic). In the case of DMIs for example, incompatibilities are expected to “snowball”, with a quadratic decline in reproductive success with genetic distance (Orr 1995; Orr and Turelli 2001). Other forms of incompatibilities might result in a linear decline, or the threshold function considered here.

**Figure 1.**
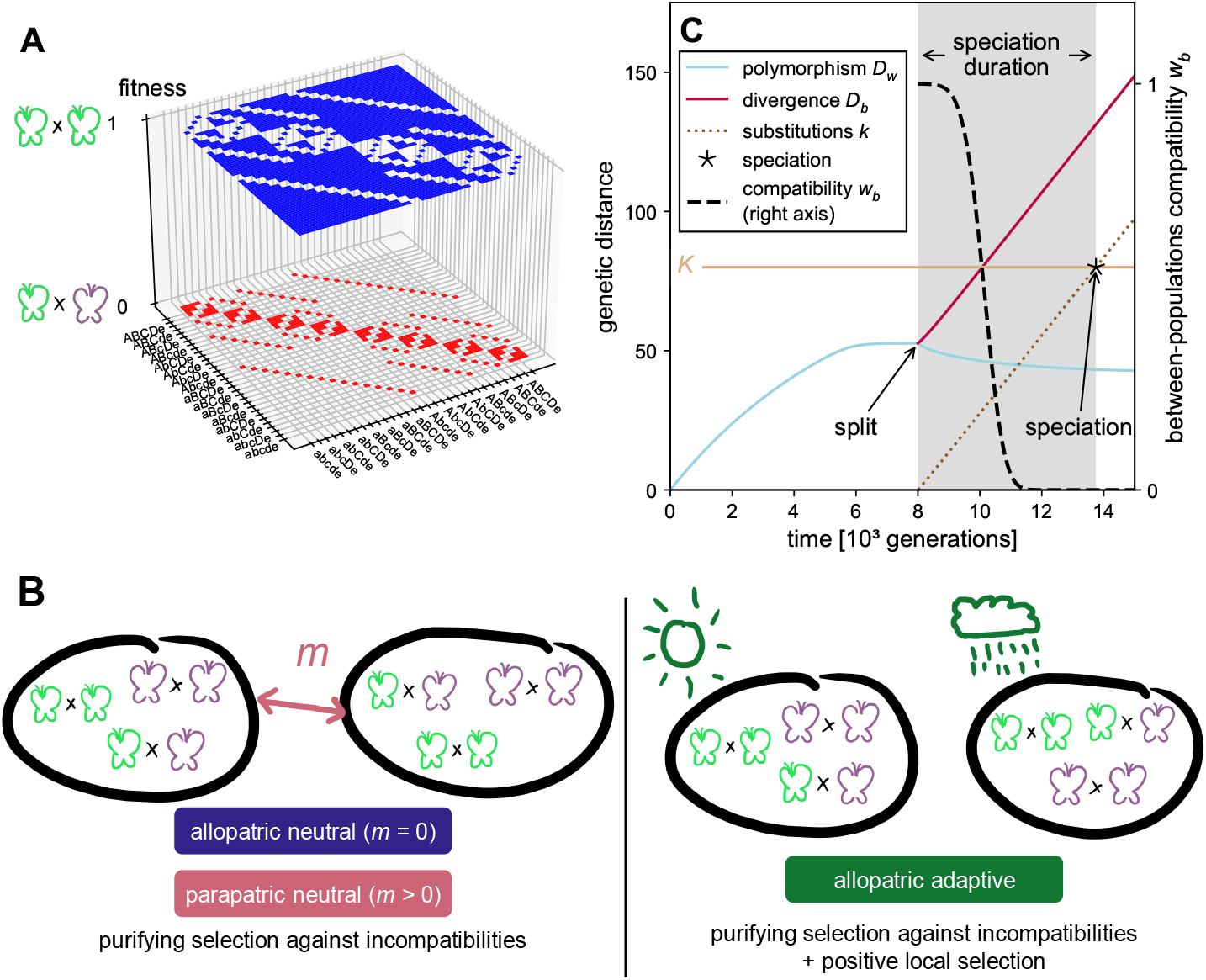
A. Illustration of the holey adaptive landscape (HAL) with 5 polymorphic sites (32 different genotypes) and a compatibility threshold *K* = 3. The genotypes have been arbitrarily sorted in alphabetical order, and only half of them are represented for readability. The fitness of genotype pairs is 1 if they differ at less than *K* sites (in blue, represented by butterflies of same color), and 0 otherwise (in red, represented by butterflies of different colors). **B.** Illustration of the different scenarios investigated in the paper: “neutral” on the left, either allopatric (in the absence of migration, *m* = 0) or parapatric (with migration, *m >* 0), and “adaptive” on the right. Purifying selection against incompatibilities operates in all scenarios, including in the neutral scenario. In the adaptive scenario, mutations in addition confer a selective advantage in the local environment in which they arise, represented by the sun and rain symbols. **C.** Example of temporal dynamics under the allopatric neutral scenario. The ancestral population is split in two after 8000 generations. We track within-population polymorphism (*D_w_*), between-population divergence (*D_b_*), the number of different substitutions between the two populations (*k*), as well as between population compatibility (*w_b_*, right axis). The star indicates speciation, characterized by the time when *w_b_* = 0, or, alternatively, *k*(*t*) = *K* with *K* the incompatibility threshold. Here *N_T_* = 6000, *N*_1_ = *N*_2_ = 3000, *ν* = 0.007, *K* = 80.

We consider three versions of the model (figure 1B). In the “allopatric neutral” scenario, the initial population experiences a random split at time *t*_split_, for example due to the emergence of a geographic barrier isolating two populations, after which strict isolation is maintained. The two populations then gradually accumulate divergence, until speciation occurs (figure 1C). By “neutral”, we mean here that mutations do not have any direct fitness effect in the local environment in which they arise; they are however not strictly neutral, as they potentially generate incompatibilities with other individuals that are selected against. In the “parapatric neutral” scenario, the two populations can exchange alleles through migration, with a migration rate *m*. In the “allopatric adaptive” scenario, mutations confer a selective advantage in the local population in which they arise, with coefficient *s*_LA_. We also discuss a potential “parapatric adaptive” scenario, which accounts for both migration and local selection. The “neutral” scenarios correspond to speciation by neutral divergence, while the “adaptive” scenarios correspond to speciation by local adaptation. We do not consider scenarios of secondary contact, where migration occurs after a prolonged period of allopatry, and therefore do not consider reinforcement mechanisms.

Following Gavrilets (1999), we use a deterministic approximation to investigate the temporal dynamics of both within-population polymorphism *D_w_*, defined as the mean pairwise distance between individuals within populations (*w* for within), and between-population divergence *D_b_*, defined as the mean pairwise distance between individuals from the two different populations (*b* for between; figure 1, Materials and methods). We also track the probabilities *w_w_* and *w_b_*that two randomly chosen individuals from the same or different populations, respectively, would successfully interbreed if they were in contact. These probabilities are referred to as within- and between-population compatibility. Finally, we follow the number *k* of substitutions (fixed alleles) that differentiate the two populations. According to the biological species concept (Mayr 1942), we consider speciation to occur if (and when) no individual from one population would successfully interbreed with any individual from the other population, should the two populations come into contact (i.e., *w_b_*= 0; Higgs and Derrida 1992; de Aguiar et al. 2009). This reproductive isolation can result from pre-zygotic barriers, post-zygotic ones, or a combination of both, and occurs if (and when) *k* reaches *K*(figure 1B); indeed, in this case, any two individuals picked in different populations are distant by at least *K* loci and are thus incompatible. We also investigate the shape of the grey zone of speciation, when the reproductive compatibility between the populations *w_b_* is reduced or even close to zero, but does not yet reach zero (Roux et al. 2016). This corresponds to the often-encountered intermediate situation in nature where populations are not fully compatible, but speciation is not complete. We validate our theoretical predictions using intensive simulations (supplementary text S2 and figures S8 to S11).

### Local adaptation accelerates speciation while gene flow delays or prevents it

In the absence of migration, populations ineluctably diverge and speciation occurs after enough time elapses. In the allopatric neutral scenario, we show that the speciation duration *τ*, defined as the expected time between *t*_split_ and the time when speciation occurs, is given by :

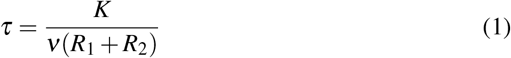

where *R*_1_ and *R*_2_ are, in each of the two sub-populations, coefficients of fixation which capture the efficiency of purifying selection to remove incompatibilities and depend on population size, mutation rate, and the incompatibility threshold (equation 3 in the Materials and methods, figure S1). For simplicity, we assume in what follows that the ancestral population is split into two populations of equal sizes, but our equations can accommodate asymmetric population sizes. Figure 2 (blue curves) shows how within-population polymorphism (lower row) and the duration of speciation (upper row) vary with *ν*, *N*, and *K*. *R* 1 represents the case when the effect of purifying selection against incompatibilities is negligible (figure 2, blue dashed curves). Equation 1 holds in the adaptive scenario, with an expression for *R* that depends on the local selective advantage of mutations *s*_LA_ (Materials and methods). As *s*_LA_ increases, the mutations in each environment confer a local adaptive advantage, and their fixation is faster. Hence the coefficient of fixation increases and can be larger than 1. As expected, local adaptation speeds up speciation (figure 2, upper row, green curves). In the presence of migration, we can determine when speciation occurs by solving a set of ordinary differential equations (Materials and methods). As expected, gene flow slows down speciation (figure 2, pink curves). Importantly, as indicated in equation 2, the accumulation of incompatibilities depends on an effective migration rate *m_e_*, which is the migration rate scaled by the between-population compatibility (higher effective migration when compatibility is higher). Hence, migration is more effective in situations that tend to slow-down the accumulation of between-population incompatibilities, rendering this accumulation even slower, and resulting in the U-shaped curves observed in the parapatric scenario. In fact, speciation is no longer ineluctable, and we designed an efficient way to predict whether it will occur or not (Materials and methods and supplementary text S1; figure 2 upper row, pink curves).

**Figure 2.**
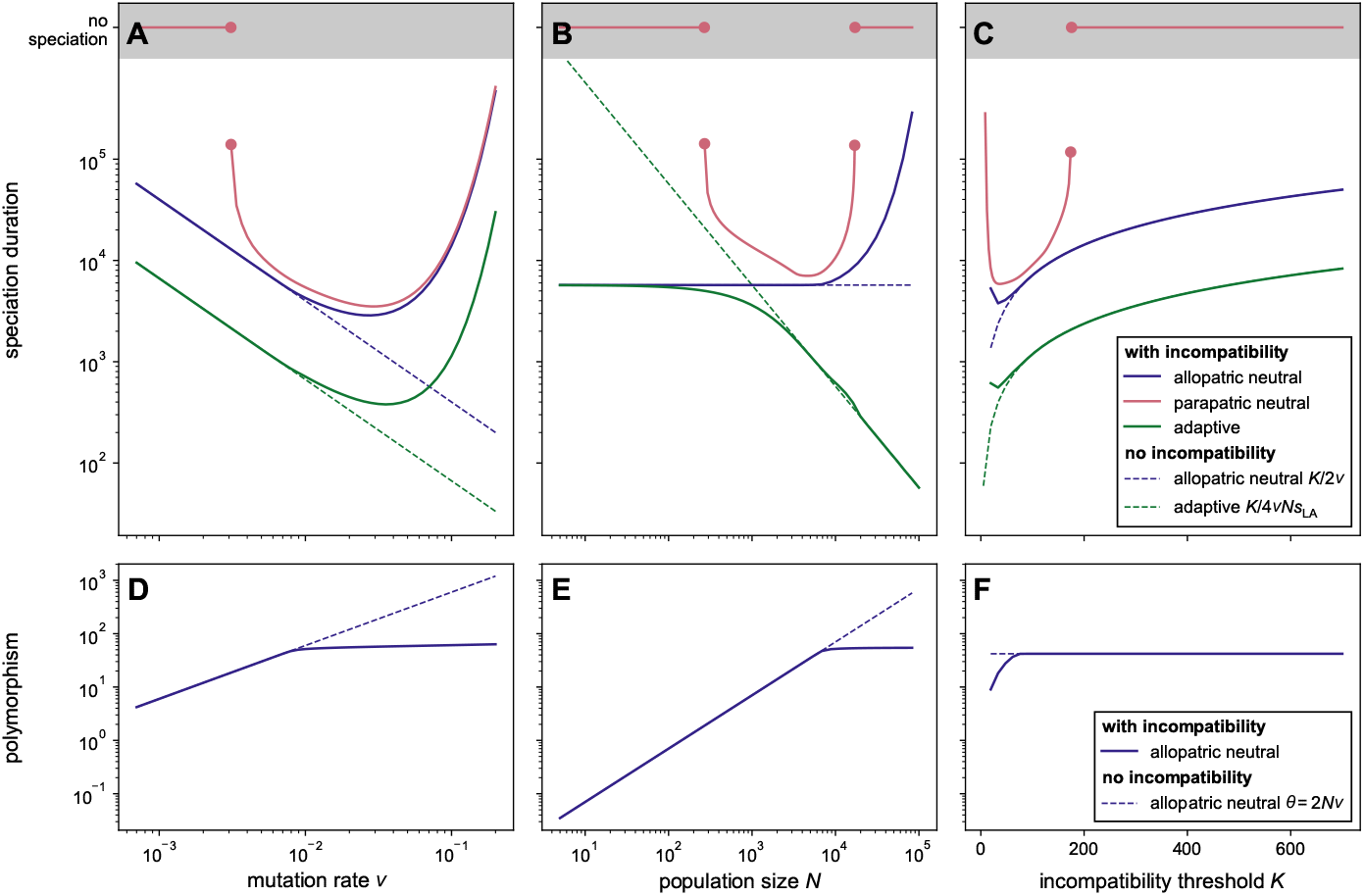
Expected speciation duration (upper row) and polymorphism (lower row) under the holey adaptive landscape model (HAL; solid lines), as a function of mutation rate (**A**, **D**), population size (**B**, **E**) and incompatibility threshold (**C**, **F**). Three versions of the model are considered: strict allopatry (in blue), with migration (parapatric, in red), and with local adaptation (in green). Expectations under a purely theoretical scenario without purifying selection against incompatibilities within populations (*R* = 1, *s* = 0) are shown with dashed lines for comparison. On the upper row, the pink line in the shaded region indicates cases where speciation does not occur. In each panel, one of the three parameters varies with the others kept constant at *ν* = 0.0007, *N* = 6000, and *K* = 80. For the parapatric scenario, the migration rate is *m* = 10*^−^*^4^ ; for the adaptive scenario, the coefficient of selection is *s*_LA_ = 0.001.

### High mutation rates can slow down speciation

High mutation rates are generally thought to increase the rate at which populations acquire substitutions and reproductive isolation, resulting in higher speciation rates (Lanfear et al. 2010; Hua and Bromham 2017). This expectation is part of the integrated evolutionary speed hypothesis, which stipulates that the short generation times and high mutation rates found in warm (tropical) environments result in fast genetic change and speciation (Rohde 1992). In a purely theoretical scenario without within-population purifying selection against incompatibilities, increased mutation rates indeed always speed up speciation (figure 2A, dashed curves). When within-population incompatibilities are selected against, however, this expectation is true only under low-to-moderate mutation rates; when mutations become more frequent (*>* 0.05 per generation), purifying selection against incompatibilities limits their accumulation both within and between populations, and slows down speciation, contrary to the evolutionary speed hypothesis (figure 2A, solid curves). Indeed, the fixation coefficient *R* decreases when the mutation rate *ν* increases (figure S1), up to a point where the fixation rate *νR* is negatively associated with *ν* (figure S2). Intuitively, this is linked to an increase in within-population polymorphism (figure 2D): in a population where individuals are already genetically distant, an individual carrying a new mutation will more likely cross the incompatibility threshold with many individuals, reducing the chance for the mutation to fix. This speciation slowdown under a high mutation rate regime occurs in the three scenarios we considered. The non-monotonic dependency of speciation duration to mutation rates may explain why tests of the evolutionary speed hypothesis have found mixed evidence. A positive association was found in birds (Lanfear et al. 2010) and plants (Bromham et al. 2015), but a negative one was found in mammals (Afonso Silva et al. 2025). In the latter study, the authors attributed this *a priori* unexpected result to methodological artifacts; our results suggest that it could in fact be real, due to a high purifying selection, and slow accumulation of incompatibilities, when mutations are frequent. In the low mutation rate regime, the increase of speciation duration when the mutation rate decreases is much stronger in the presence of migration than in strict allopatry (figure 2A, pink curve). Intuitively, mutations not only get rarer, they are also less likely to fix as *ν* gets smaller, because between-population divergence is lower, implying that migrant genotypes are more likely to be compatible with residents, and to homogenize populations (i.e., migration is more effective).

### Faster speciation in smaller populations is a signature of neutral divergence

The relationship between speciation duration and population size strongly depends on whether genetic divergence is driven by locally advantageous mutations (adaptive scenario, in green) or occurs without positive selection on the mutations contributing to divergence (neutral scenarios, in blue and red) (figure 2B). As expected, speciation by local adaptation is faster in large populations where positive selection is more effective; this effect becomes visible when the population size is larger than the drift-selection barrier (1*/*4*s*_LA_ = 250 here; Kimura and Ohta 1971). This prediction would also hold in the presence of migration (whether the mutations that confer a local selective advantage are neutral or deleterious in the other population), as migrants, which are less fit than individuals from the local population, will be more rapidly selected against in large populations, reducing their chance of homogenizing diverging populations. In the allopatric neutral scenario, and in a purely theoretical situation without within-population purifying selection against incompatibilities, population size would not have an effect on the duration of speciation (figure 2B, dashed blue curve). When within-population incompatibilities are selected against, however, this purifying selection is more efficient in larger populations, slowing down both the accumulation of incompatibilities and speciation (figure 2B & E, solid blue curve). After a given size threshold, the relationship between population size and speciation duration is thus positive. The same holds with migration, except that, when populations decrease below a certain size threshold, speciation duration increases as populations become smaller (figure 2B, pink curve). Intuitively, between-population divergence — which is equal to within-population polymorphism when populations split — is lower in small populations (panel E), making migration all the more effective and homogenizing. Importantly, these results show that faster speciation in larger populations can occur under both adaptive and neutral divergence, but faster speciation in smaller populations occurs only under speciation by neutral divergence (with or without gene flow). We additionally find that speciation duration is primarily influenced by the size of the smallest of the two separate populations in speciation by neutral divergence, whereas it is primarily influenced by the size of the largest one in speciation by local adaptation (figure S5). This makes intuitive sense, as mutations causing reproductive incompatibilities accumulate faster in the smallest of the two populations if they are not adaptive, and in the largest population if they are adaptive. We do not see any reason why these results would change with the form of the relationship between genetic distance and reproductive success, although this would need to be directly investigated.

### Speciation is slower when reproductive isolation is more polygenic

The incompatibility threshold *K*, i.e., the number of genetic differences at which individuals no longer interbreed, reflects the genomic architecture of speciation, ranging from few large-effect genes (or structural genomic changes) for small *K* to many small-effect genes spread across the genome for large *K*. These genomic changes can result in pre- and/or postzygotic barriers, which have the same consequence of leading to the reproductive isolation of genetically distant individuals. In a purely theoretical scenario without within-population purifying selection against incompatibilities, the duration of speciation increases with *K*, as a larger number of fixed differences is required for speciation to occur (figure 2C, dashed curves). It is also largely the case when incompatibilities are selected against, except for very small values of *K*, where speciation duration increases (figure 2C, solid curves). This is due to the extreme purifying selection acting on genomic changes with large effects, which hampers their fixation (see the reduced coefficient of fixation *R* in figure S1 for small *K*) and slows down speciation, despite the small number of fixed differences required for speciation to occur; this effect is reflected by a reduced polymorphism in the population (figure 2F). Hence, the more polygenic speciation is, the more time it is expected to take, except in cases of very large effect genomic changes that have a hard time fixing. The presence of migration amplifies the effect of *K* on speciation duration (figure 2C, pink curve). When *K* is small, this can be attributed to the decline of within-population polymorphism, and therefore also of between-population divergence, which renders migration more efficient. When *K* is large, increasing *K* not only increases the between-population divergence threshold at which speciation occurs, but also the chance that migrants are compatible with residents, and therefore homogenize populations.

### Gene flow extends the grey zone of speciation

Mutation rates, population sizes, and the genomic architecture of speciation influence not only speciation duration, but also the timing and shape of the grey zone (GZ), i.e., when and at which pace the between-population compatibility declines. We fitted a decreasing sigmoid to the between-population compatibility *w_b_*(*t*) (dashed line in figure 1B) and analyzed its timing *t*_GZ_ (inflection point) and slope *r*_GZ_ (Materials and methods). The results mirror the dependencies found for speciation duration (see the concordance between figure 3A and figure 2A–C, see also figure S3), with earlier/faster drops in between-population compatibility corresponding to shorter speciation durations (e.g., in the presence of local adaptation when population sizes are large), while later/slower drops correspond to longer speciation durations (e.g., in the presence of migration). The grey zone can also reveal effects that are not captured by speciation duration: in the case of small populations and in the absence of migration, the duration of speciation does not change with population size (figure 2B, on the left), but the slope of the grey zone does, with slower compatibility drops when population size increases (figure 3A2, vertical part of the blue and green curves). The only scenario with an early drop in between-population compatibility is the scenario with local adaptation when populations are large (figure 3A2, in green), and migration enforces a slow compatibility drop (figure 3A2, in red). Hence, we expect — most often — a substantial lag between the time when populations split and the time when compatibility between them drops, and fuzzy species boundaries that extend for a long period of time as soon as there is gene flow between-populations.

**Figure 3.**
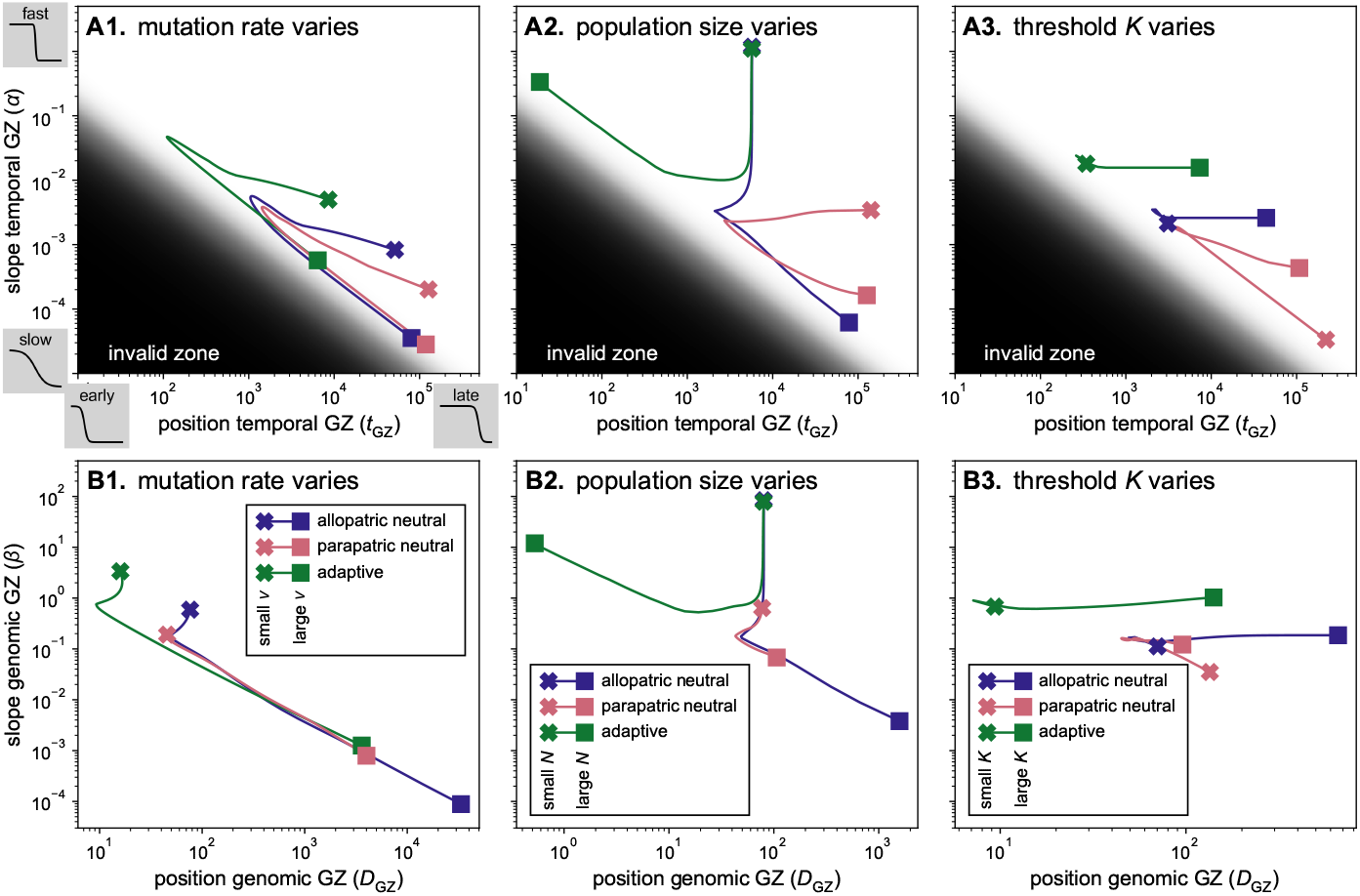
Characteristics of the temporal (**A**) and genomic (**B**) grey zone (GZ) of speciation, with varying mutation rate (**1**), population size (**2**) and incompatibility threshold (**3**). The temporal GZ (resp. genomic GZ) is characterized by a slope *α* (resp. *β* ) and a position *t*_GZ_ (resp. *D*_GZ_), see Materials and methods. Each line represents how the slope and position of the GZ change when one parameter changes (increasing from ✖ to ▪), under a specific scenario of speciation. On the upper row, the dark zone of the graphics indicates mathematically unreachable regions; typically early speciation with small slope would require a low initial compatibility *w_b_*(the middle of the border correspond to *w_b_* = 0.75). The values of both the fixed and varying parameters are the same as in figure 2.

### Interpreting empirical grey zones of speciation

We also examined the shape of the grey zone as a function of the net synonymous divergence (defined as the difference between divergence and polymorphism at synonymous sites, *D*^syno^ := *D*^syno^ *− D*^syno^) instead of time, as is typically done in genomic-based empirical analyses of the grey zone of speciation (Materials and methods; Roux et al. 2016; Monnet et al. 2025). At constant mutation rate (figure 3B and C), the genomic grey zone behaves similarly as the temporal grey zone, except in the scenario with migration. Indeed, at constant mutation rate in the absence of migration, net synonymous divergence is a good proxy for time. With migration, net divergence no longer increases linearly with time, and variations in population size and threshold *K* have a small effect on the position and slope of the genomic grey zone, although they have a strong effect on the duration of speciation. These analyses further show that, when speciation occurs, the presence of migration does not affect the position or slope of the genomic grey zone in comparison with the case without migration. Indeed, migration reduces both the accumulation of incompatibilities and the between-population net synonymous divergence at a comparable pace, such that the effects cancel one another. The most drastic case of incongruence between the temporal and genomic grey zones occurs when mutation rates vary, in the low mutation rate regime. In this regime, speciation is faster at higher mutation rates, but the genomic grey zone gives the opposite impression. Its slope becomes flatter when mutation rates increase, but this reflects the faster increase of synonymous divergence rather than a deceleration of speciation.

Our theoretical results on the genomic grey zone should help interpreting the shape of the grey zone obtained from empirical data (Roux et al. 2016; Monnet et al. 2025). For example, Monnet et al. (2025) found that reproductive isolation is achieved at a smaller level of divergence in plants compared to animals. They hypothesized that this difference could be explained by reproductive isolation occurring with fewer reproductive barriers in plants (for example, Moyle and Graham 2005). We show here that the number of loci required to achieve reproductive isolation (*K*) indeed has a large effect on the position of the grey zone (figure 3B). Our results suggest that differences in population sizes, smaller mutation rates, and/or stronger effects of local adaptation in plants compared to animals could also explain the pattern. Differences in gene flow, to the contrary, are not expected to impact the shape of the genomic grey zone. When comparing genomic grey zones in empirical systems, a difference in slope could be due to differences in mutation rates, population sizes, and strength of local selection, but not in the number of reproductive barriers.

### Genomic data in plants reveal faster speciation in smaller populations, indicative of neutral divergence

Our theoretical predictions indicated that a positive correlation between population size and the duration of speciation would be indicative of speciation by neutral divergence and that, in this case, speciation duration should be primarily influenced by the size of the smallest population. We evaluated these correlations using genomic-based demographic inferences for 196 pairs of plant species for which speciation was inferred to be complete (i.e., no gene flow at present; Monnet et al. 2025). We used posterior distribution estimates of speciation duration and of the sizes of the ancestral and two descendant populations, and compared the correlations with a null model to account for the structure in pairs of the data (see subsection 4.4). The correlation between speciation duration and population size was significantly positive (*p <* 0.05) in 98.8% of the posterior samples when considering the small descendant population (25% in the null model), 59.5% of the samples when considering the larger descendant population (22.2% in the null model), and none of the samples when considering the ancestral population (22% in the null model; figure 4). The absence of correlation between the ancestral population size and the duration of speciation may seem surprising at first sight, since population size typically influences the initial polymorphism within- and between-populations. However, the positive correlation between speciation duration and descendant population sizes suggests that we are in the large-population regime (figure 2B, right-most part) where polymorphism is expected to be independent of population size (figure 2E, right-most part). The empirical correlations are therefore consistent with expectations under speciation by neutral divergence and large population sizes.

**Figure 4.**
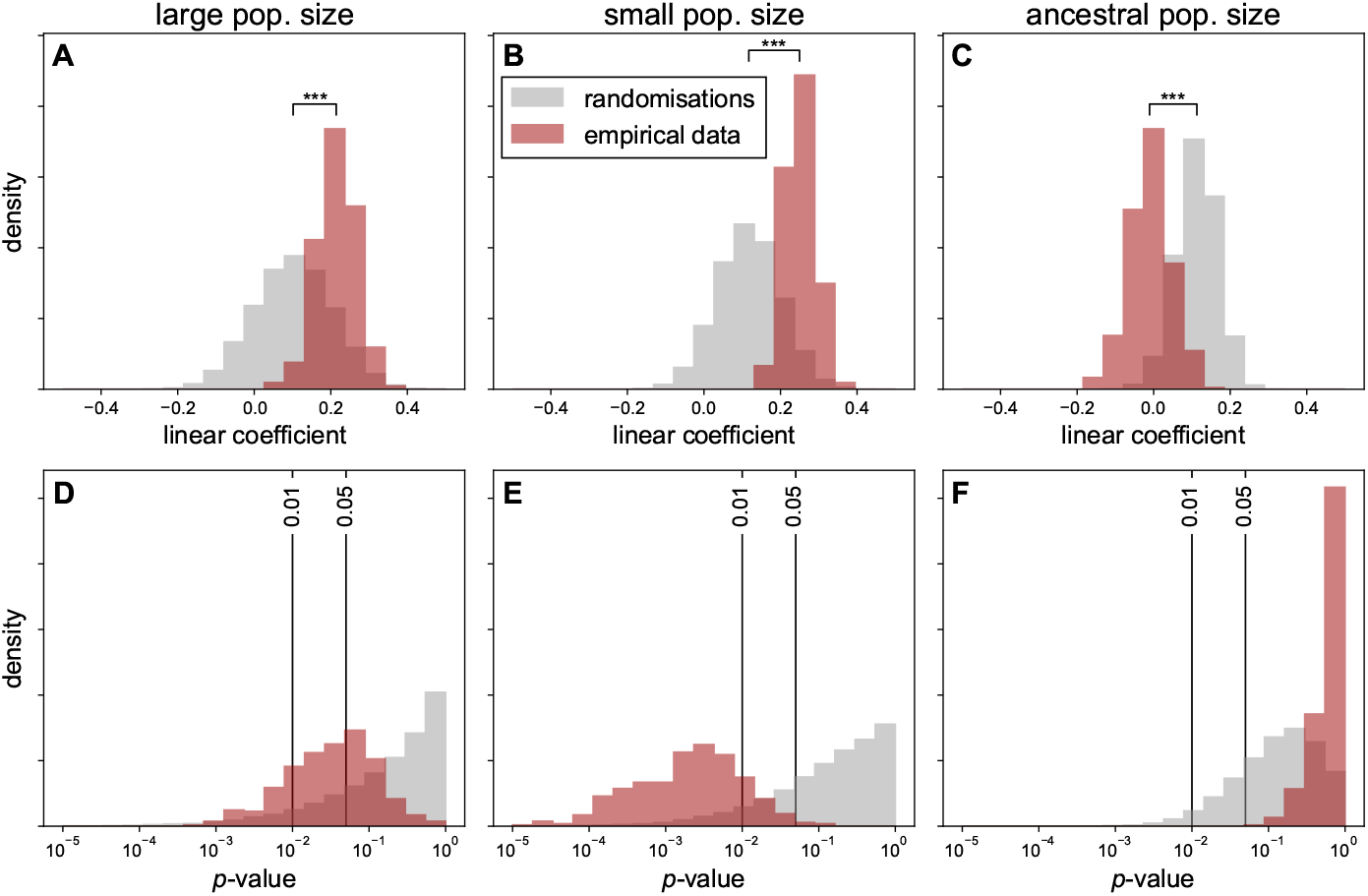
Distribution of linear coefficients (**A**, **B**, **C**) and *p*-values (**D**, **E**, **F**) for the correlation between speciation duration and population size across posterior samples (red) and randomisations (grey). We fitted the model: log(spec. time) log *N*_large_ + log *N*_small_ +log *N*_ancestral_ to 400 posterior samples from demographic parameters inferred on 196 pairs of plants, compared with 100 randomisations between pairs of species for each sample. On the upper row, the difference between the linear coefficient from the randomisations and the empirical data is assessed with a paired *t*-test (see detailed results in table S1).

The consistency of the empirical correlation between population size and speciation duration with expectations under a non-adaptive speciation scenario does not contradict years of empirical evidence for local adaptation in plants (Fournier-Level et al. 2011; Lowry 2012). Instead, it suggests a decoupling between the traits under local adaptation and the genetic differences causing reproductive isolation (Frayer et al. 2025), such that non-adaptive rather than adaptive processes control the duration of speciation. Under this scenario, as incompatibilities accumulate between populations, less gene flow will counteract local adaptation, and ecological differences between sister species will arise more easily (van der Niet and Johnson 2009). The ecological differences would thus be facilitated by reproductive isolation rather than serving as its initial cause (Bierne et al. 2011). An alternative explanation for the observed positive correlation between speciation duration and population size is that factors promoting the accumulation of incompatibilities are stronger in small populations, such as selection and geographical isolation. For example, small populations may frequently originate from long-distance dispersal into novel habitats, leading to increased geographic isolation. Upon arrival in markedly different environments, they would also experience stronger, more divergent selection. If small populations generally arose as a result of such long-distance dispersal, we would expect less gene flow in smaller populations. We did not find any significant relationship between ancestral population size and ancestral migration rates estimated from the plant genomic data (figure S12); however, this could simply be due to the difficulty of inferring past migration rates (Fraïsse et al. 2021). Regardless of the mechanisms at play, our results suggest that one should not necessarily expect speciation to be faster in larger populations.

## 3 Conclusion

We investigated how mutation rates, population sizes, the genomic architecture of speciation, selection, and migration shape the tempo of speciation in a multigenic, correlated adaptive landscape. As expected, higher mutation rates accelerated speciation, consistent with the evolutionary speed hypothesis, whereas speciation proceeded more slowly when mutations were neutral than when they were locally adaptive and in the presence of gene flow. We also uncovered several unexpected patterns, including a reversal of the relationship between mutation rates and speciation duration expected under the evolutionary speed hypothesis at very high mutation rates, and the insensitivity of the genomic grey zone of speciation to gene flow.

Our theoretical and empirical results linking speciation duration to population size revive Mayr’s verbal theory of genetic revolution. Specifically, speciation can occur rapidly in small populations in the absence of local adaptation, due to the increased possibility of fixing incompatibilities, and empirical data from 196 plant species pairs are consistent with this non-adaptive speciation scenario. In a widely cited perspective, Schluter (2009) opened by noting that “evolutionary biologists (…) finally agree that the origin of species occurs by means of natural selection”, by which he meant positive selection to the environment, during ecological or mutation-order speciation. Our results challenge this view by suggesting that the origin of species may be driven primarily by the accumulation and fixation through genetic drift of incompatibilities subject to purifying selection, rather than by positive selection. Nearly neutral evolution may therefore not only dominate molecular evolution but also be a major determinant of the tempo of species formation.

If smaller populations tend to speciate faster, they are also more vulnerable to extinction and may collapse before speciation is complete. This trade-off underscores the need, in the future, to jointly consider speciation duration and lineage persistence to understand how microevolutionary processes translate into macroevolutionary speciation rates. The framework developed here provides a foundation for the development of such integrative analyses.

## 4 Materials and methods

### 4.1 Predictions under the holey adaptive landscape model

#### Neutral scenario: results from Gavrilets (1999)

Following Gavrilets (1999), we use a system of ordinary differential equations (ODE) to approximate the mean polymorphism *D_w_* (genetic distance between pairs of individuals in a population), the mean divergence *D_b_* (genetic distance between pairs of individuals from two separate populations), and the number of distinct substitutions (fixed alleles) between the two populations *k*. Under the assumptions of linkage equilibrium and rare alleles, the within- and between populations mean compatibilities are given by (Gavrilets 1999, eq. 12 and 15):

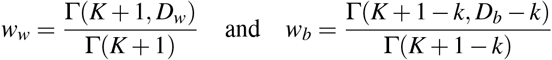

with Γ( , ) and Γ( ) the upper incomplete gamma function and complete gamma function, respectively. The effect of purifying selection against incompatibilities is captured by a selection coefficient that depends on the level of within-population polymorphism (Gavrilets 1999, eq. 11):

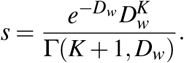

In the absence of local adaptation, the dynamics follow the system of equations:

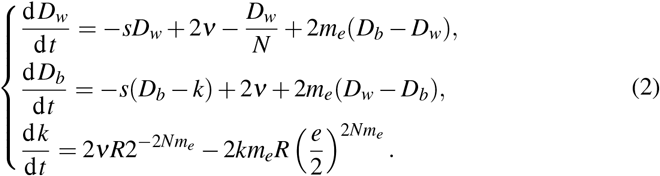

where *N* is the size of each population, *ν* is the number of mutations per individual per generation, denoted thereafter as mutation rate, *m_e_* := *m w_b_/w_w_* is the effective migration rate, *m* the migration rate (probability for an individual to have migrated from the other population at the previous generation), *s* is the selection coefficient defined above, and *R* a fixation coefficient capturing the relative speed of allele fixation compared to a hypothetical case without incompatibility. The expression of *R* is given by (Gavrilets 1999, eq. 13b):

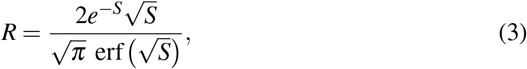

with erf the error function and *S* := *Ns/*2.

#### Neutral scenario: original results

In the particular case without migration (*m* = *m_e_*= 0), and under the assumption that *D_w_* reached an equilibrium (so *R* is constant), we can obtain an analytical solution for the duration of speciation by solving the equation for *k*, which reduces to 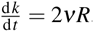. When the ancestral population splits, individuals are distributed randomly in the two populations, and it is unlikely that an allele is absent in one population and fixed in the other. Therefore, if *t* = 0 represents the splitting time, the initial number of distinct substitutions is *k*(0) = 0, and thus *k*(*t*) = 2*νRt*. Considering that speciation is achieved when *k*(*t*) = *K*, the duration of speciation is given by 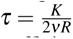. If the two populations have different sizes *N*_1_ and *N*_2_, and therefore different coefficients of fixation *R*_1_ and *R*_2_, the equation becomes equation 1.

To analyse the dynamics in the neutral scenarios, we numerically solved the system given in equation 2. In the presence of migration, speciation is not systematic. In this case, predicting whether speciation will occur or not based on the ODEs can be challenging and time consuming. We designed an alternative, fast approach, based on the bifurcation diagram for the equilibriums of this system. The detailed method is provided in supplementary text S1.

#### Scenario with local adaptation: results from Gavrilets (1999)

To analyse the dynamics in the allopatric scenario with local adaptation, we used the system of ODEs:

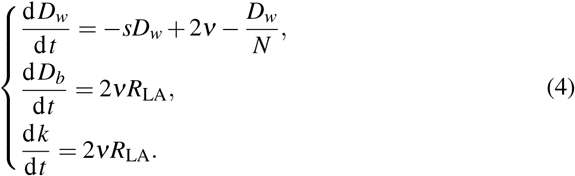

with *R*_LA_ the coefficient of fixation in the case of local adaptation, given by (Gavrilets 1999, eq 19):

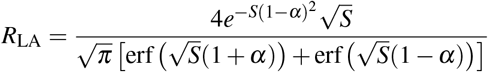

with *α* = *s*_LA_*/s* denoting the importance of selection due to adaptation compared to the selection due to incompatibility. The equation for *D_w_* is an approximation that ignores the effect of local adaptation (the equation is the same as in the neutral scenario with *m_e_*= 0) and is justified by the fact that increases in polymorphism due to positive selection when alleles are in low frequency are counter-balanced by decreases when they are in high frequency. The equation for *D_b_* is an asymptotic approximation (Gavrilets 1999, eq. 13a); this equation shows that *R*_LA_ captures the acceleration of between-population genetic divergence induced by the divergent selection in the two populations. In this model, the mutations responsible for incompatibilities have a selective advantage in the population in which they appear. In this allopatric model the two populations do not exchange any gene, such that the selective status of mutations in the other population (i.e., neutral, deleterious or advantageous) is irrelevant.

#### Implementation

We used Python 3.13 and the module scipy (Virtanen et al. 2020) to numerically solve the ODEs. We provide a tool to resolve the equations directly in command line, available on github.com/pierre-veron/HoleyAdaptSpeciation. In order to speed up convergence, we used some numerical approximations for some of the calculations, such as the fixation coefficient. Our implementation provides the option to use either the exact form or the approximation.

### 4.2 Simulations

The predictions above rely on several simplifications: rare allele assumption, linkage equilibrium, approximation of stochastic processes by deterministic equations, and additional approximations in the scenario with local adaptation. To verify the validity of these simplifications, we compared the output of the equations with stochastic simulations (supplementary text S2). We provide a tool to run the simulations in command line, available on github.com/pierre-veron/HoleyAdaptSpeciation. Without migration, we find that the equations predict well the dynamics observed under the simulations, in particular when we added recombination to the simulations to approximate the linkage equilibrium assumption (figures S8 to S11, configurations A2 and B2). In the neutral allopatric scenario, even simulations without recombination matched the predictions well (condition A1). In the presence of migration to the contrary (conditions C to H), outcomes are more stochastic, and some simulations lead to speciation while it is not expected based on our equations, especially when the migration rate is high and populations are small (*N* = 100 for each population) (conditions G and H). This is consistent with the deterministic approximation, which is not valid when populations are small.

### 4.3 Analyses of the grey zone of speciation

The grey zone (GZ) of speciation is the period before speciation during which the two populations are not fully compatible nor fully incompatible. We characterized the GZ by analyzing the between-population compatibility *w_b_* as a function of either *(i)* time (the temporal GZ) or *(ii)* net synonymous divergence (the genomic GZ). The net synonmymous divergence is defined as *D_a_*^syno^ := *D_b_*^syno^ *− D_w_*^syno^, where *D_w_*^syno^ and *D_b_*syno are the polymorphism and divergence on sites where there is no selection for incompatibility (see supplementary text S3 for the equations). This analysis of the compatibility as a function of the net synonymous divergence is similar to empirical studies on genomic data (e.g., Roux et al. 2016; Monnet et al. 2025). We fitted a decreasing sigmoid to these curves. The temporal GZ is thus characterized by *w_b_*(*t*) = 1*/*(1 + *e^α^*^(*t*^*^−t^*^GZ)^), with *α* the slope, and *t*_GZ_ the timing, or position, of the temporal GZ. The genomic GZ is similarly characterized by *w* (*D_a_*^syno^) = 1*/*(1 + *e^β^*^(*D*^*_a_*^syno^*^−D^*_GZ_ ^)^) with *β* the slope, and *D*_GZ_ the position, of the GZ. We performed the fit using the function curve fit from the package scipy and we checked that the sigmoid was a good approximation (coefficient of determination *R*^2^ *>* 0.971). We investigated how the slope and position of the GZ vary with mutation rates, population size and threshold of incompatibility, and between the three scenarios of speciation.

### 4.4 Empirical link between population size and the duration of speciation

We used genomic estimates of various population genetic parameters from Monnet et al. (2025). These were obtained by applying the Demographic Inference with Linked Selection (DILS) model (Fraïsse et al. 2021) to 280 pairs of plant species (118 species or populations from 25 genera). DILS outputs a best supported model among ancient migration (AM, recovered in 196 pairs), secondary contact (SC, 46 pairs) and isolation with migration (IM, 38 pairs); given our goal to retrieve estimates of speciation duration, we selected the 196 species pairs for which speciation was inferred to be complete, i.e., the AM model (see figure S4). We retrieved posterior distributions of the time of the split (*T*_split_) and the time of cessation of gene flow (*T*_AM_), and used their difference as our estimate of speciation duration (*T*_spec_). We also retrieved the posterior distributions of the sizes of the ancestral (*N_a_*) and two descendant populations (*N*_large_ for the largest, and *N*_small_ for the smallest). We tested the relationship between the duration of speciation and the population sizes by fitting an ordinary least square (OLS) regression: log *T*_spec_ log *N*_large_ + log *N*_small_ + log *N_a_* and repeated this over 400 samples.

The species pairs we used are possible pairs within each of 25 plant genera (Monnet et al. 2025). Individual species therefore appear in multiple pairs, generating a complex pattern of non-independence in the data, structured per genus. To account for this non-independence, we repeated the analysis after randomly permuting species identities within each genus. A potential correlation obtained for these permuted data cannot be explained by a true correlation with population sizes, but is rather due to repeated patterns in the data structure (see figure S4). Comparisons between empirical correlations and null model correlations (i.e., those obtained after random permutations) allow for the detection of “true correlations” not due to the way the data are organised. For each empirical sampled, we fitted an OLS on 100 randomised data and compared the obtained linear coefficients using a paired *t*-test (see table S1).

## 5 Aknowledgements

We thank Sergey Gavrilets, Camille Roux, Quentin Rougemont, Nicolas Bierne and Emmanuel Schertzer for their help with the model or data. PV acknowledges funding from the *École polytechnique* and the *Institut des mathématiques pour la Planète Terre* (https://impt.math.cnrs.fr/).

## 6 Data availability

All scripts and data used in this analysis are available on Zenodo: https://doi.org/10.5281/zenodo.18683264. A GitHub repository containing the functions to run HAL predictions and simulations in command line is also available: https://github.com/pierre-veron/HoleyAdaptSpeciation/.

---

**figure S1.**
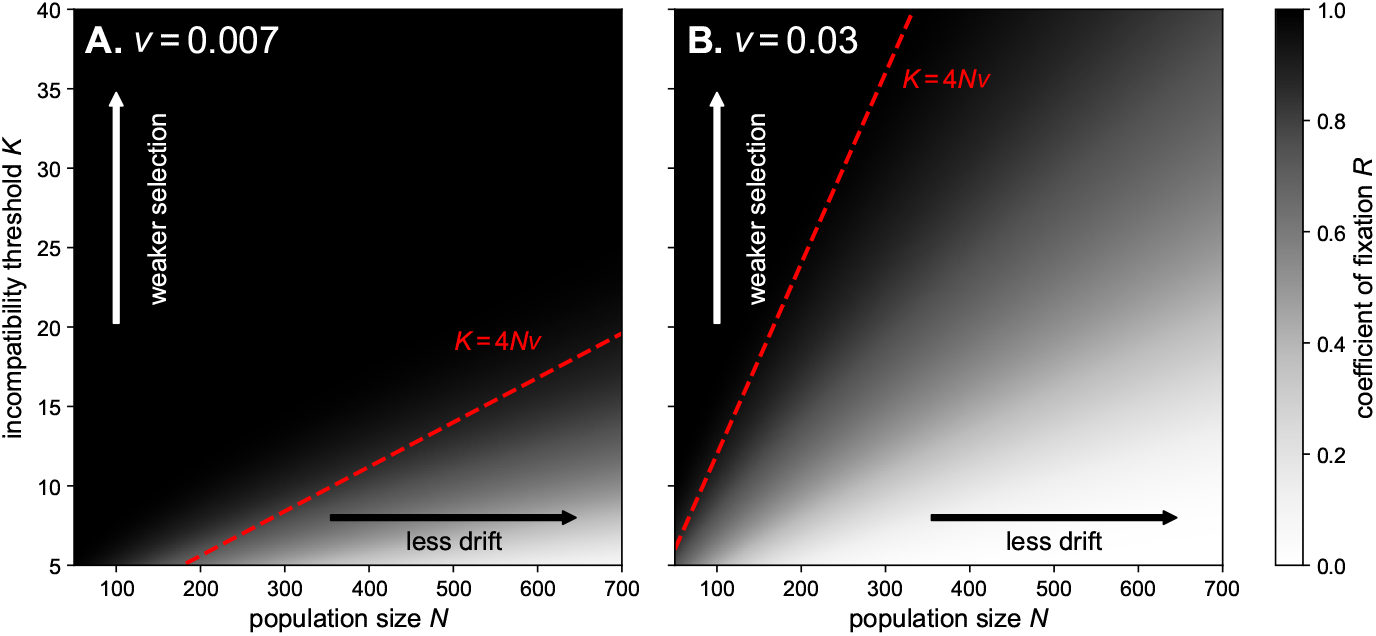
Values of the coefficient of fixation *R* (representing the relative speed of fixation of mutations compared to a case without purifying selection against incompatibilities, see equation 3) as a function of the population size *N* and the incompatibility threshold *K*, for two values of mutation rate *ν* (**A** and **B**). The dotted red line shows the rule of thumb proposed by Gavrilets (1999) for the limiting conditions where purifying selection against incompatibilities can be neglected (*R ≈* 1).

**figure S2.**
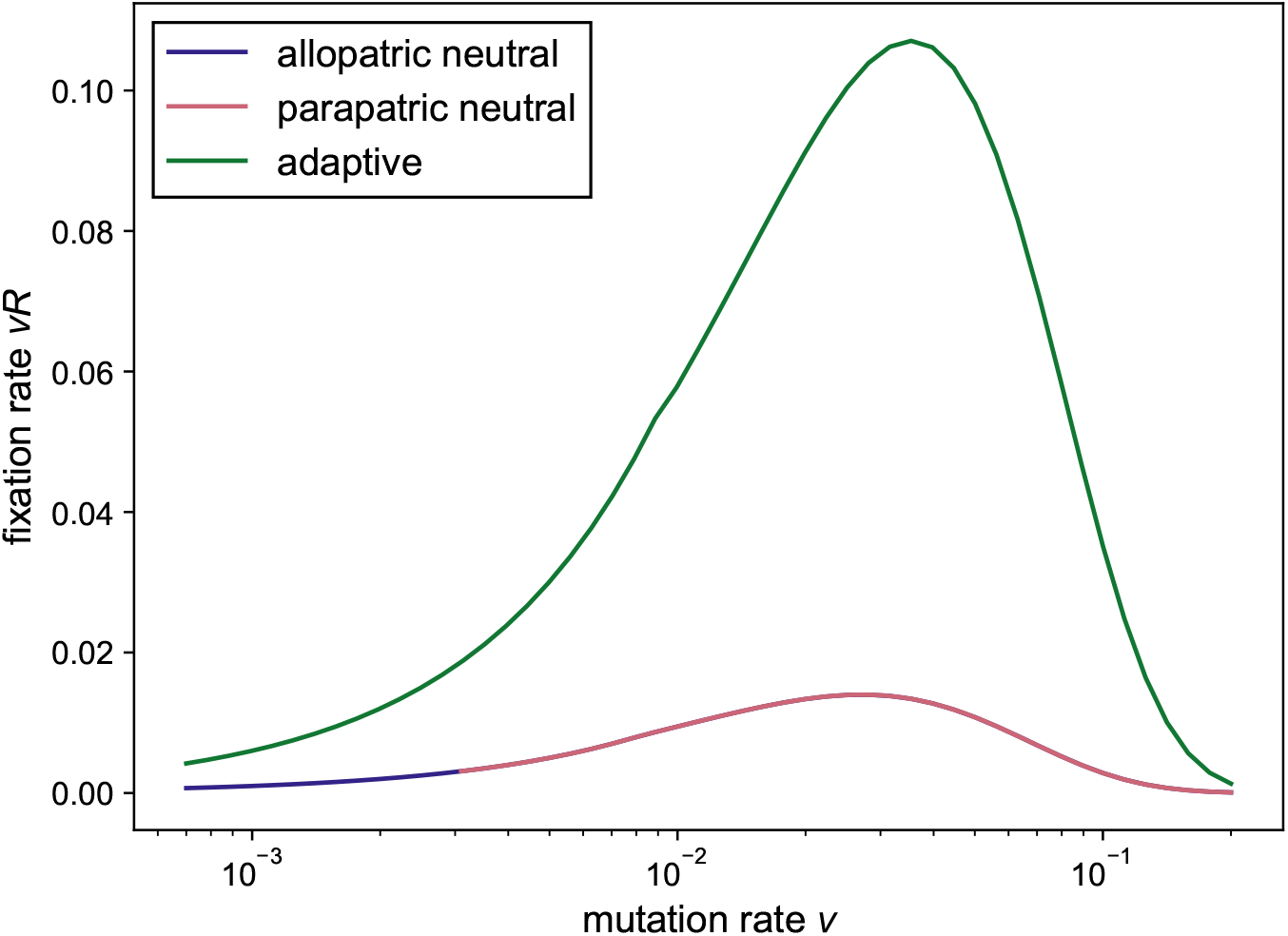
Fixation rate *νR* as a function of the mutation rate *ν* in the three scenarios.

**figure S3.**
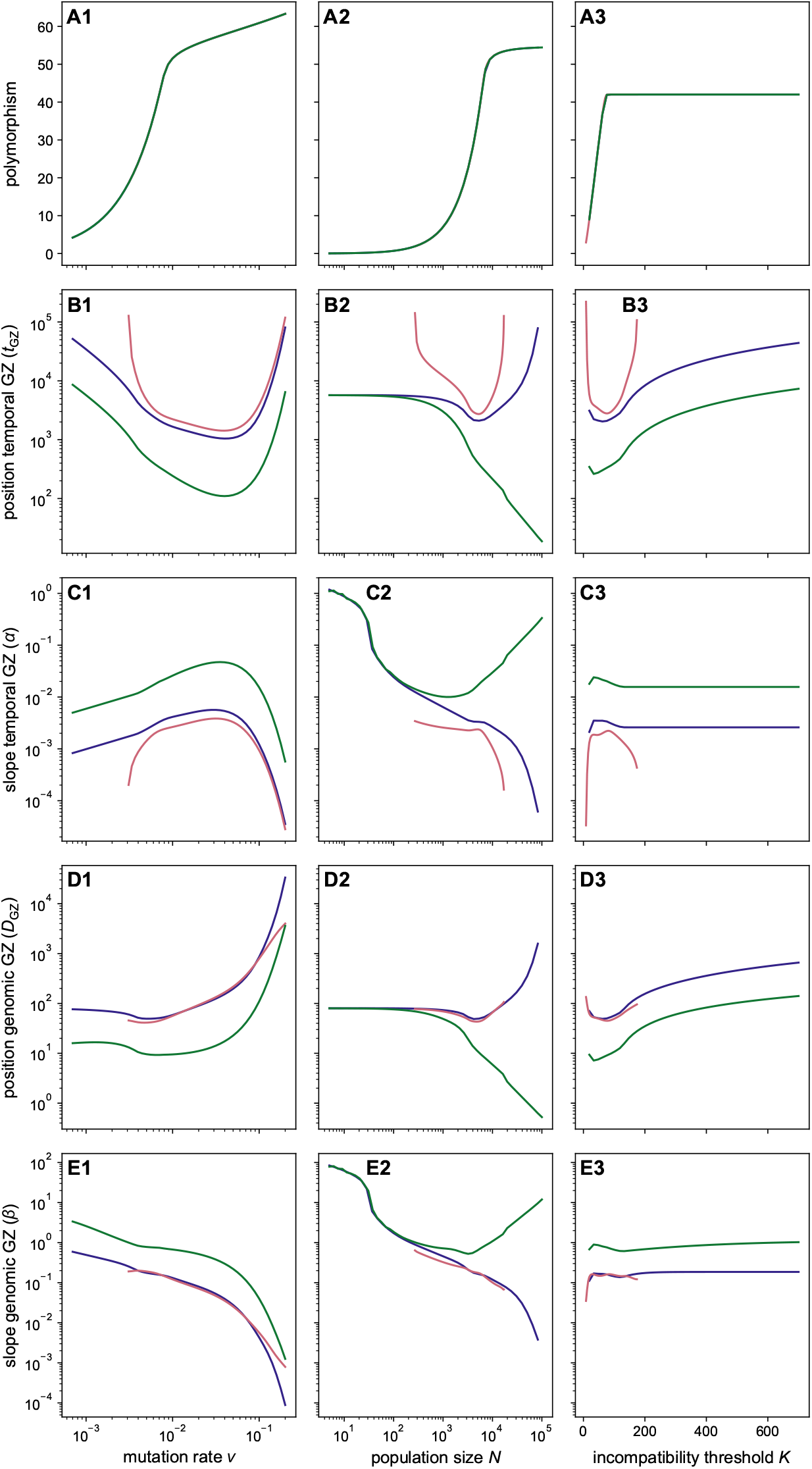
Details of the characteristics of the temporal and genomic grey zones (GZ) of speciation as a function of the mutation rate (**1**), the population size (**2**) and the incompatibility threshold (**3**). **A** shows the mean polymorphism, **B** and **C** show the position and slope of the temporal GZ and **D** and **E** show the position and the slope of the genomic GZ. Each line corresponds to a scenario of speciation.

**figure S4.**
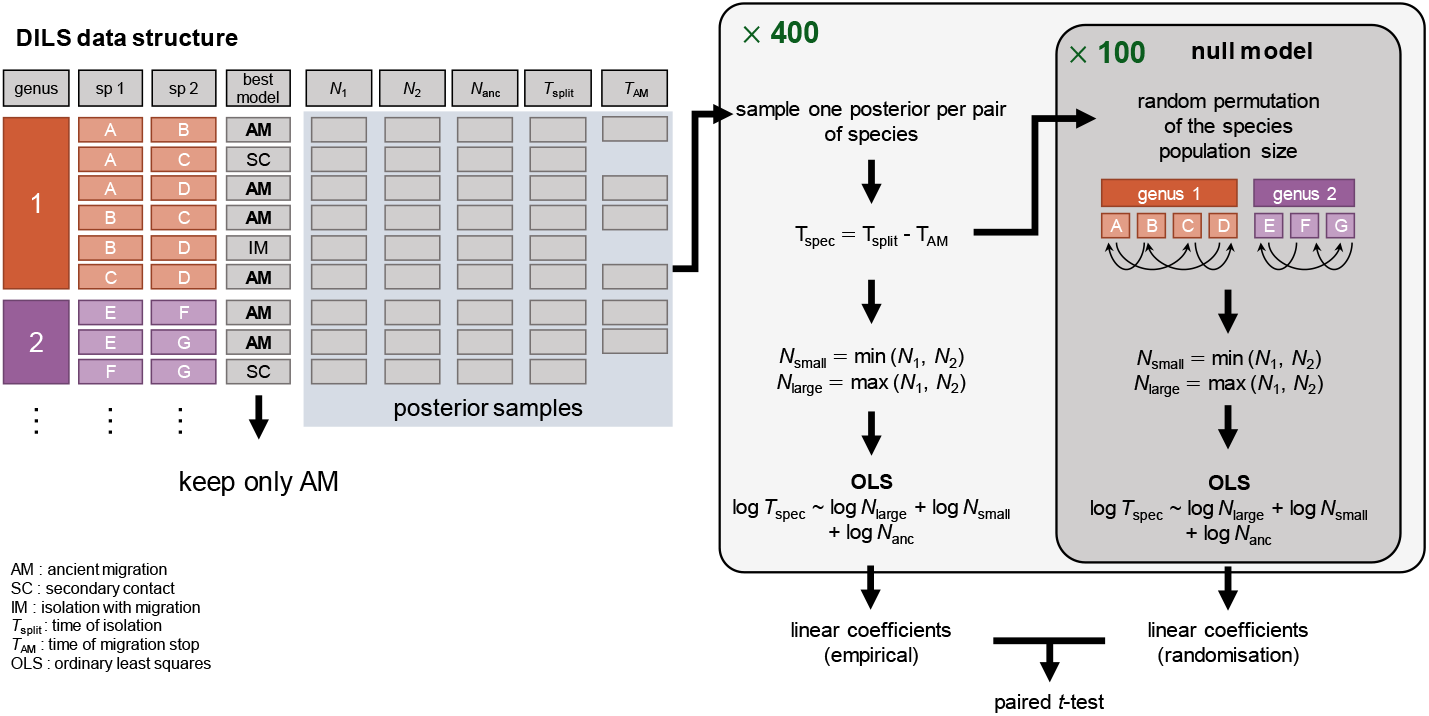
Illustration of the paired structure of the data obtained from DILS, and of the statistical approach used to test the significance of the regression between speciation duration and population sizes . We permuted species within each plant genus to construct a null model.

**figure S5.**
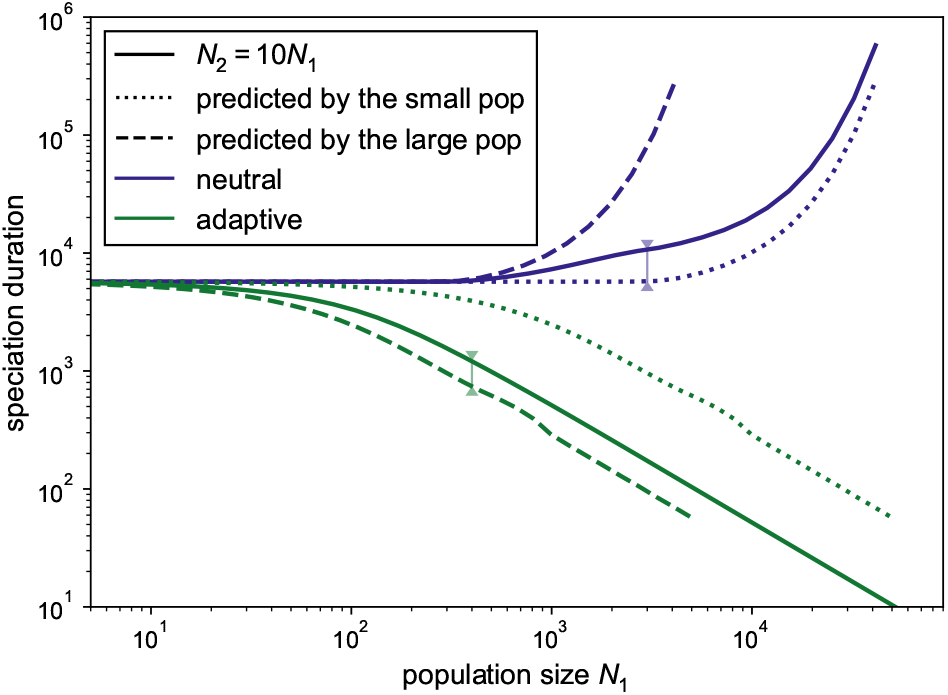
Duration of speciation under the holey adaptive landscape (HAL) allopatric models, in the case of asymmetric populations (solid lines), as a function of *N*_1_ with *N*_2_ = 10*N*_1_, compared to the prediction with two populations with size *N*_1_ (dotted line) and two populations with size *N*_2_ (dashed line). In the absence of local adaptation, speciation duration is primarily driven and best predicted by the size of the smallest population (see the small difference between the solid and dotted blue lines, blue arrow); in the presence of local adaptation, it is instead best predicted by the size of the largest population (green arrow).

**figure S6.**
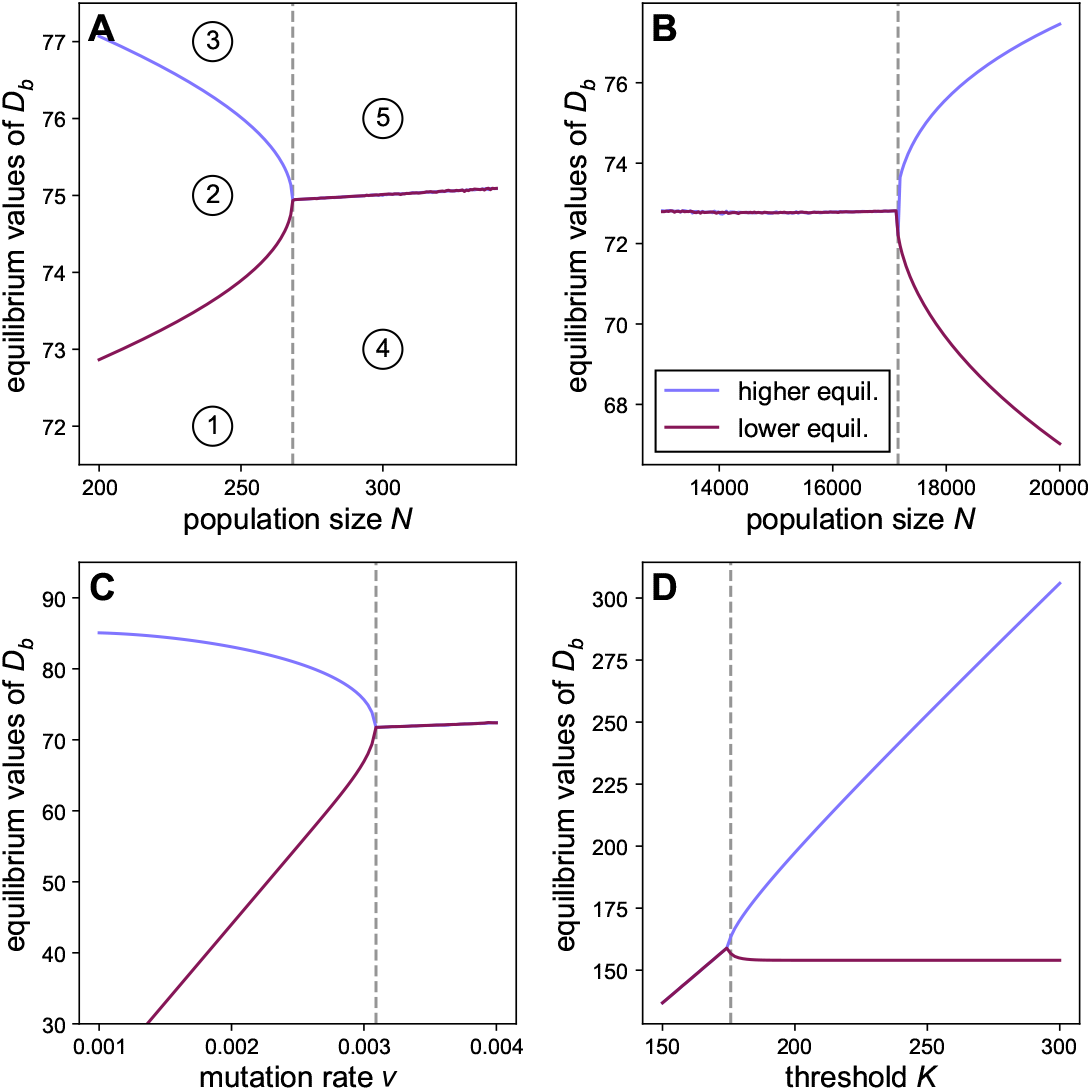
Bifurcation diagrams showing the equilibrium value(s) for *D_b_* as a function of the population size (**A**: low values, **B**: high values), the mutation rate *ν* (**C**) and the threshold of incompatibility *K* (**D**). When one parameter varies, the others are kept constant: *ν* = 0.0007, *N* = 6000, *m* = 10*^−^*^4^ and *K* = 80. The numbers on panel **A** indicate initial conditions for which the dynamics of *D_b_* are represented on figure S7.

**figure S7.**
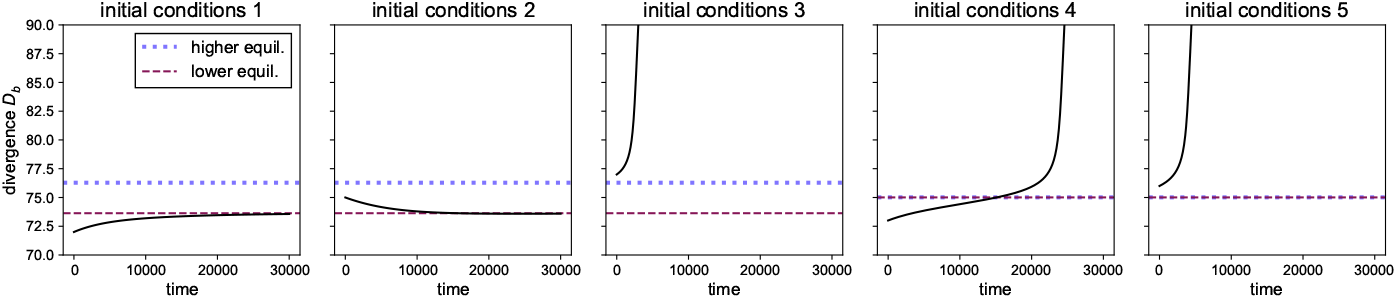
Dynamics of *D_b_* when its initial value is close to the equilibrium and smaller than (1, 4), higher than (3, 5), or between (2) the equilibrium values, corresponding to 5 different regions of the bifurcation diagrams, indicated on figure S6A. The dotted lines indicate the equilibrium value(s). In each case the initial values of *D_w_*and *k* are the equilibrium values.

**figure S8.**
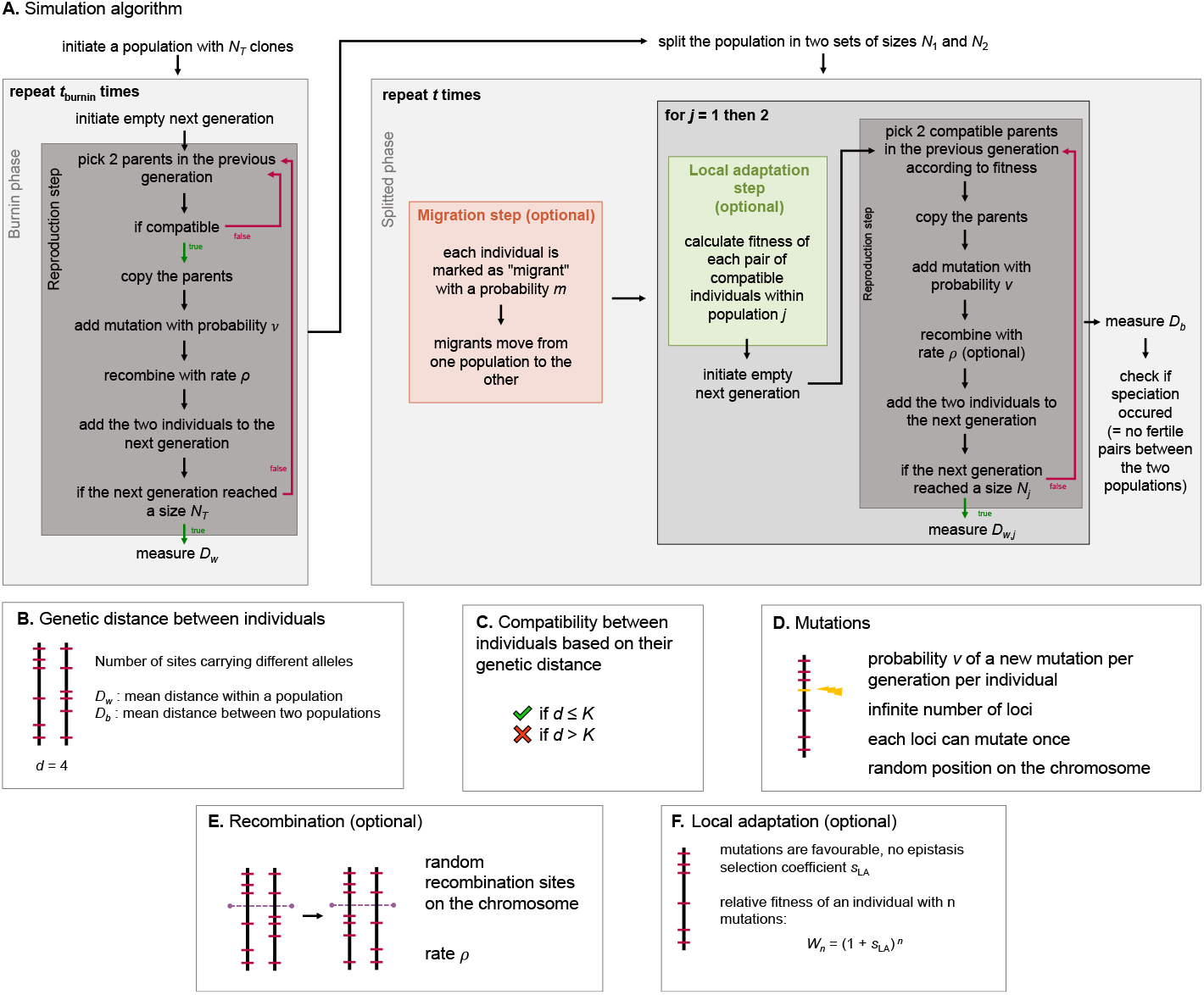
Illustration of the simulation algorithm (**A**) and the several components of the process (**B** to **F**). **A**: A burnin phase is used to allow the ancestral population to reach an equilibrium polymorphism. After this phase, the populations are splitted randomly . **B**: The distance between individuals is calculated as the number of sites carrying different alleles. **C**: The HAL model assumes that individuals can reproduce if their genetic distance is smaller than the incompatibility threshold. **D**: At each generation, each individual can mutate with probability *µ*. In this case, a site that never carried a mutation is altered. A gene cannot undergo the same mutation twice independently. **E**: For simulations with recombination, a part of the chromosome is randomly exchanged between the two offsprings of a pair of parents. **F**: For simulations with local adaptation, the fitness of the each compatible pair of individuals within each populations is calculated and is used to sample the parents during the reproduction step.

**figure S9.**
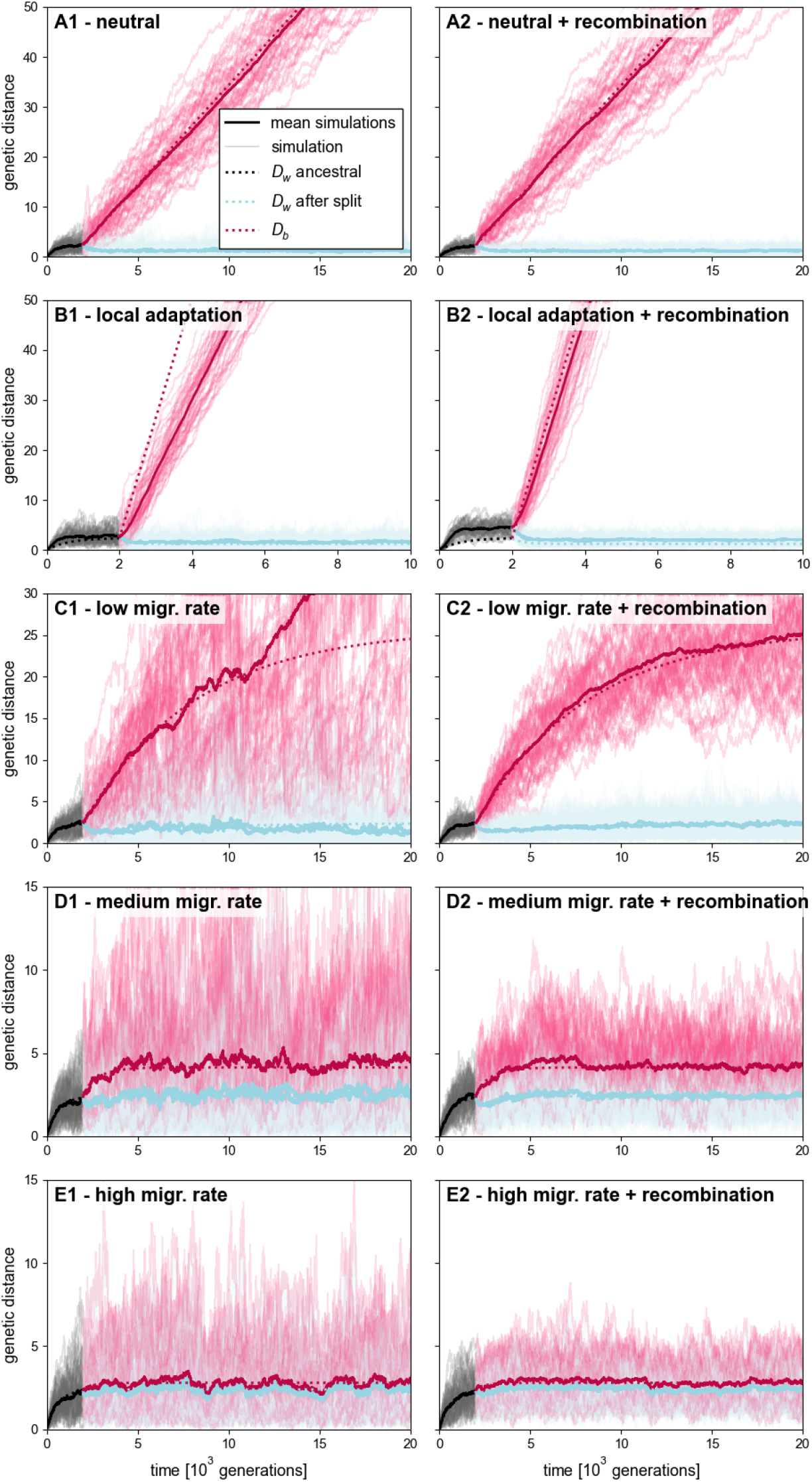
Dynamics of polymorphism and divergence under the holey adaptive landscape model obtained with the stochastic simulations (50 simulations for each configuration, solid thin lines ; the solid thick lines show the mean of the simulations) and with the deterministic approximations (dotted lines). The first column shows simulations without recombination and the second column shows simulations with recombination (recombination rate *ρ* = 2). The first row (**A**) corresponds to the allopatric neutral model, the second row (**B**) corresponds to the allopatric model with local adaptation (*s*_LA_ = 0.005) and the last third rows (**C**, **D**, **E**) correspond to the model with low, medium and high migration (migration rate *m* = 8.5 10*^−^*^5^, 0.001155 and 0.00498 respectively). Parameters of the model: *ν* = 0.002*, N*_1_ = *N*_2_ = 300*, K* = 40, burnin time 2000 generations.

**figure S10.**
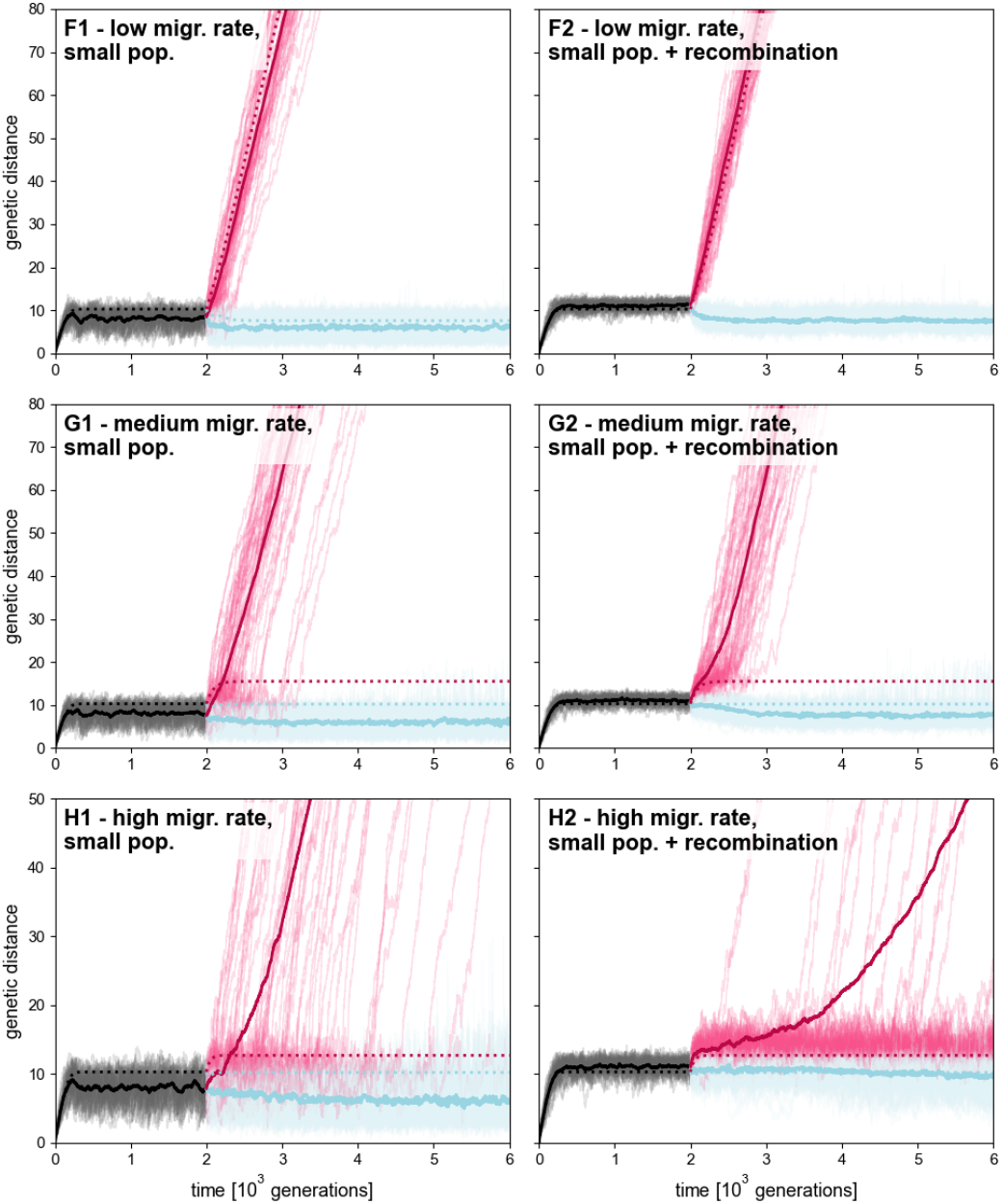
Same as figure S9 but with smaller populations : *ν* = 0.0384, *N*_1_ = *N*_2_ = 100, *K* = 20 and migrations rates 0.001, 0.005 and 0.01 for rows **F**, **G** and **H** respectively.

**figure S11.**
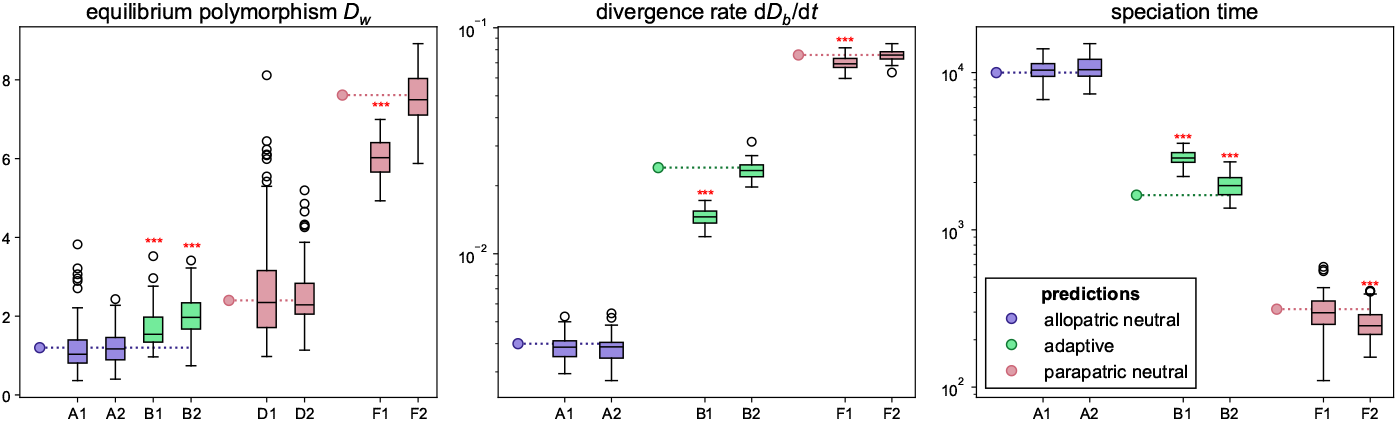
Comparison of the dynamics of the simulations and the predictions, based on (left) the equilibrium polymorphism *D_w_*, (middle) the slope of *D_b_*(*t*) and (right) the duration of speciation. The boxplots indicate the distribution of the estimates from 50 simulations for each configuration, and the dot the expected value based on the deterministic prediction. The configurations (A1, A2, B1, B2, D1, D2, F1 and F2) are those of figures S9 and S10. The red asterisks indicate significant differences, assessed with *t*-test and corrected for multiple testing.

**figure S12.**
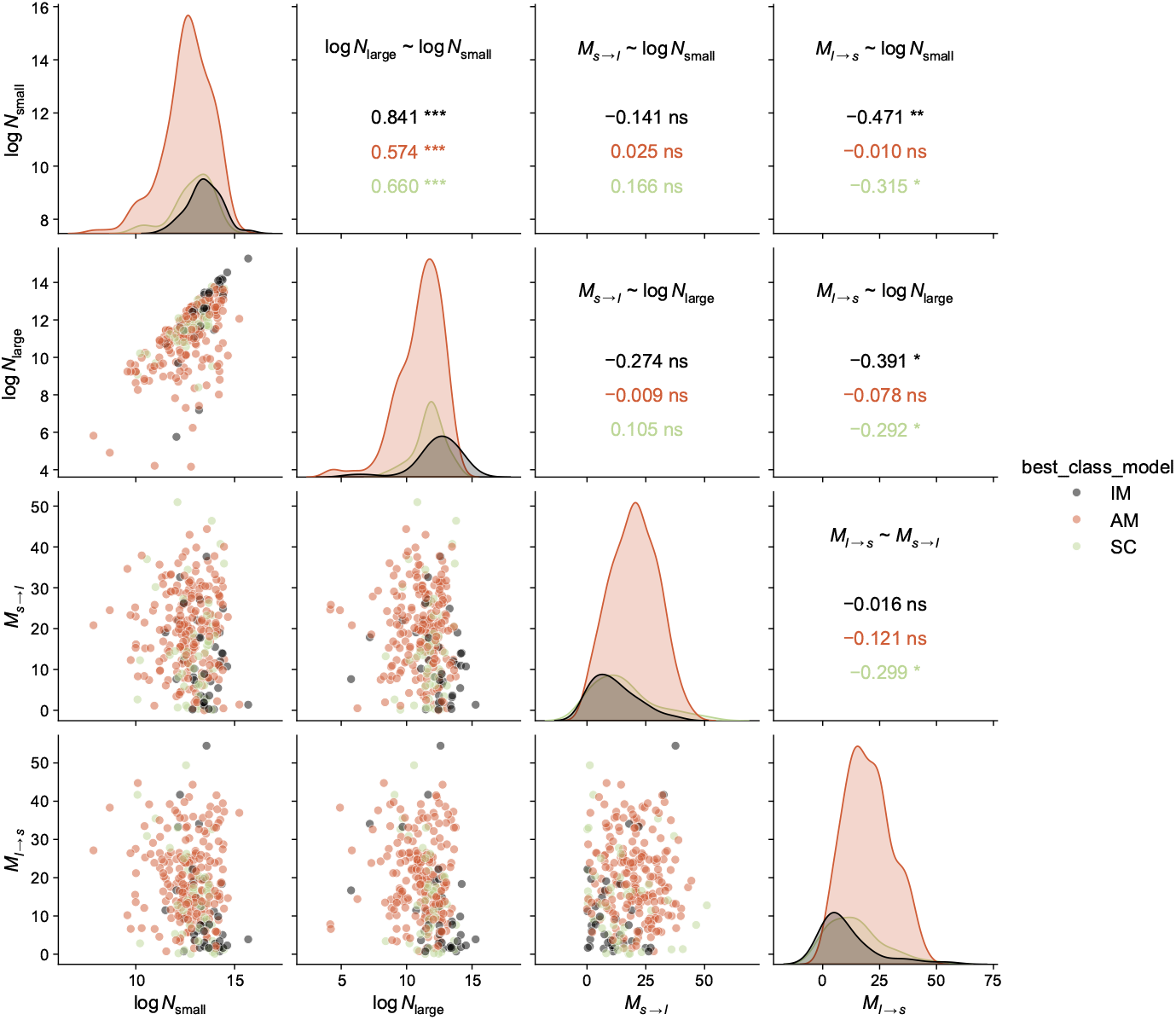
Correlation between average log-population size and migration rates inferred by DILS in the data by Monnet et al. (2025). *M_s→l_* correspond to the average number of migrants going from the small population to the large population at each generation. The upper corner indicates Spearman’s correlation coefficient.

**Table S1.**
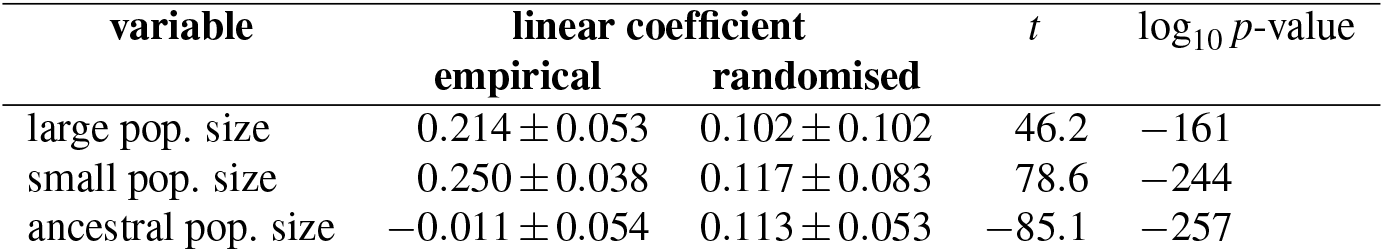
Details of the paired *t*-test between the linear coefficient of the OLS regression on the empirical versus randomized data (*n* = 400 samples).

**Supplementary text S1.** Determination of the cases without speciation under the parapatric model

In the presence of migration, there are combinations of parameters for which speciation does not occur. Determining these cases by solving the ODEs in equation 2 is not straightforward and time consuming (in particular, speciation not occurring within the set time frame does not imply it will never occur). We therefore designed an alternative approach based on the analyses of the equilibria of the differential equations. For a given combination of parameters, we computed the equilibria by numerically finding the values *D_b_*, *D_w_*and *k* such that the time derivatives provided in equation 2 are zero. Depending on the parameter values, there are either two equilibrium conditions (one with large *D_b_*, and one with small *D_b_*) or one equilibrium (the two values coincide, see the bifurcation diagrams on figure S6). These equilibrium conditions are not necessarily met, as they require a specific combination of polymorphism, divergence and number of substitutions. The dynamics of *D_b_*(*t*) around these equilibria determine whether speciation occurs or not, with *D_b_*(*t*) diverging to ∞ when speciation occurs. We tested numerically the stability of the equilibrium conditions by resolving *D_b_*(*t*) with an initial condition close to the equilibrium value and *D_w_*, *k* taken equal to the equilibrium values. In the cases with one equilibrium, we found that this equilibrium is unstable (*D_b_* ∞ as *t* ∞; figure S7 cases 4 and 5) ; in the cases with two equilibria (figure S7 cases 1, 2 and 3), the lower equilibrium is stable (*D_b_* converges to this equilibrium) and the higher equilibrium is unstable (*D_b_* ∞). In practice, the initial divergence *D_b_*(0) is small compared to the equilibrium values, such that condition 3 is not met. Hence, in the parameter space with two equilibria, *D_b_* converges to the lower, stable equilibrium and speciation does not occur. In the parameter space with one equilibrium, to the contrary, speciation occurs and *D_b_*increases without limit. The bifurcation points shown on figure S6 (dotted lines) can thus be used to determine the parameter values at which there is a transition between scenarios with or without speciation (as in figure 2).

**Supplementary text S2.** Comparison of the HAL deterministic model with simulations

The equations we used to investigate speciation under the HAL model rely on several simplifications, in particular a deterministic approximation (stochastic processes are modelled by ordinary differential equations), an assumption of linkage equilibrium (the allele frequency at a locus is assumed to be independent of the allele frequency at the other loci), a rare allele approximation (the frequency of alleles at polymorphic sites is assumed to be close to 0 or 1 most of the time), and additional approximations detailed in Materials and methods in the case with local adaptation. In reality, the stochasticity around the equilibrium can play a role in the duration of speciation, especially in small populations where the random reproduction of a small number of individuals can have a strong effect on the dynamics of the whole population. This is exacerbated in the case with migration, as the few individuals who migrate may or may not be compatible with individuals from the receiving population, with important consequences for the speciation process. Furthermore, the assumption of linkage equilibrium is not necessarily met, for example if the recombination rate is low. Gavrilets (1999) performed simulations without recombination and interpreted observed discrepancies between predictions and simulations as the effect of linked loci. We implemented here stochastic individual-based simulations that can account for recombination. To assess the conditions under which the equations are valid and the effect of the linkage equilibrium assumption, we compared the outcomes of the equations with simulations including (or not) recombination. Our simulation process is detailed in figure S8. figure S9 and figure S10 report the results of the comparison with “intermediate” (*N* = 600) and small (*N* = 200) population sizes, which are then summarized in figure S11. With larger population sizes, we expect predictions to be closer to simulations as the deterministic approximation is more valid.

**Supplementary text S3.** Dynamics of the synonymous polymorphism and divergence

For the purpose of measuring the genomic grey zone of speciation, we calculated the synonymous polymorphism *D_w_*^syno^ and the synonymous divergence *D_b_*^syno^, alongside the non-synonymous ones. Synonymous polymorphism and divergence correspond to the genetic diversity that would accumulate on regions of the genome where there would be no selection, neither due to adaptation or incompatibility.

In a single population of size *N*, the synonymous polymorphism follows the equation:

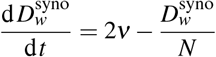

Therefore, *D_w_*^syno^ reaches the equilibrium *D_w_*^syno^*^,∗^* = 2*νN*. We assume that the polymorphism and divergence on synonymous sites at the time when populations split is given by this equilibrium. Next, they follow the system of equations:

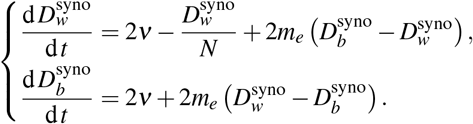

These equations were obtained by setting the selection coefficient *s* of equation 2 to 0, so that the only forces driving diversity are mutation, genetic drift and migration. Although we consider synonymous sites under no selection here, the incompatibilities accumulating on the other sites modulate this last force, which is expressed by the effective migration rate *m_e_*.

